# The Early Diagnosis in Lung Cancer by the Detection of Circulating Tumor DNA

**DOI:** 10.1101/189340

**Authors:** Geng Tian, Xiaohua Li, Yuancai Xie, Feiyue Xu, Dan Yu, Fengjun Cao, Xuanbin Wang, Fenglei Yu, Weiquan Zhong, Shixin Lu, Xiaonian Tu, Xumei Yao, Jiankui He, Chaoyu Liu

**Affiliations:** Department of Oncology, Shenzhen Second People’s Hospital, Shenzhen, China; Shenzhen GeneHealth Bio Tech Co., Ltd., Shenzhen, Guangdong, 518053, China; Thoracic Department, Peking University Shenzhen Hospital, Shenzhen, China; Laboratory of Chinese Herbal Pharmacology, Oncology Center, Renmin Hospital, Hubei University of Medicine, Shiyan, 442000, China; Hubei Key Laboratory of Wudang Local Chinese Medicine Research, Shiyan, 442000, China; Department of thoracic surgery, The Second Xiangya Hospital of Central South University, Changsha, China; Department of thoracic surgery, Huizhou Third People’s Hospital of Guangzhou Medical University, Huizhou, China; Department of Biology, South University of Science and Technology of China, Shenzhen, 518055, China

## Abstract

Remarkable advances for clinical diagnosis and treatment in cancers including lung cancer involve cell-free circulating tumor DNA (ctDNA) detection through next generation sequencing. However, before the sensitivity and specificity of ctDNA detection can be widely recognized, the consistency of mutations in tumor tissue and ctDNA should be evaluated. The urgency of this consistency is extremely obvious in lung cancer to which great attention has been paid to in liquid biopsy field.

**Methods:** We have developed an approach named systematic error correction sequencing (Sec-Seq) to improve the evaluation of sequence alterations in circulating cell-free DNA. Averagely 10 ml preoperative blood samples were collected from 30 patients containing pulmonary space occupying pathological changes by traditional clinic diagnosis. cfDNA from plasma, genomic DNA from white blood cells, and genomic DNA from solid tumor of above patients were extracted and constructed as libraries for each sample before subjected to sequencing by a panel contains 50 cancer-associated genes encompassing 29 kb by custom probe hybridization capture with average depth >40000, 7000, or 6300 folds respectively.

**Results:** Detection limit for mutant allele frequency in our study was 0.1%. The sequencing results were analyzed by bioinformatic expertise based on our previous studies on the baseline mutation profiling of circulating cell-free DNA and the clinicopathological data of these patients. Among all the lung cancer patients, 78% patients were predicted as positive by ctDNA sequencing when the shreshold was defined as at least one of the hotspot mutations detected in the blood (ctDNA) was also detected in tumor tissue. Pneumonia and pulmonary tuberculosis were detected as negative according to the above standard. When evaluating all hotspots in driver genes in the panel, 24% mutations detected in tumor tissue (tDNA) were also detected in patients blood (ctDNA). When evaluating all genetic variations in the panel, including all the driver genes and passenger genes, 28% detected in tumor tissue (tDNA) were also detected in patients blood (ctDNA). Positive detection rates of plasma ctDNA in stage I lung cancer patients is 85%, compared with 17% of tumor biomarkers.

**Conclusion:** We demonstrated the importance of sequencing both circulating cell-free DNA and genomic DNA in tumor tissue for ctDNA detection in lung cancer currently. We also determined and confirmed the consistency of ctDNA and tumor tissue through NGS according to the criteria explored in our studies. Our strategy can initially distinguish the lung cancer from benign lesions of lung. Our work shows that the consistency will be benefited from the optimization in sensitivity and specificity in ctDNA detection.

## Introduction

A growing number of newly diagnosed cancers at the advanced stages are recognized worldwide every year^1,2^, and methods and tools enabling earlier diagnosis are extremely urgent to improve the curative treatment and life quality for cancer patients^3^. Biomarkers consisting of circulating tumor cells, circulating free nucleic acids and exosomes are reported to carry the information of tumorigenesis and provide the clue for early detection of cancers. With the progress of next-generation sequencing (NGS) in recent years, new technology for the detection of these biomarkers called “liquid biopsy” is being developed and attracting more and more attention. Liquid biopsy is expected to substitute solid biopsy in the long term for its noninvasion and convenience in the diagnosis, medication, recurrence monitoring and prognosis assessment of cancers. It is therefore believed to play critical roles in precision medicine and personalized medicine of cancers^4,5^.

Among all the substrates of blood-based liquid biopsy, circulating free DNA (cfDNA) in the blood circulation system has emerged as the most important biomarker because of its availability and stability compared with circulating tumor cells, circulating free RNA and exosomes. The cfDNA in the plasma of cancer patients has been believed to contain circulating tumor DNA (ctDNA) for decades, although over ninety percent of cfDNA in healthy individuals is from metabolism of normal blood cells. Previous studies suggested that cfDNA can serve as a sensitive tool for early diagnosis of cancers^6-10^ and the levels of ctDNA were reported to increase with the severity of cancers^11,12^. Currently, ctDNA is gradually recognized as a significant biomarker for cancer monitoring and treatmen^t3,5,^ especially after several research groups reported the positive detection of ctDNA in early-stagy cancers^13-16^.

However, there are still several difficulties remained to overcome before ctDNA be can widely accepted and used in direct detection of early cancers. Firstly, the amount of ctDNA is limited in blood. The fact that only 0.01%-0.1% of the plasma cfDNA are tumor-related ctDNA and the truth that currently the highest recovery rate of ctDNA is around 10-20% mean some early cancers are not detectable due to inhesion factor, so technique for ctDNA extraction and enrichment with higher recovery must be developed. Secondly, baseline reference is needed. The removal of the background somatic mutations (the noise) and non-cancer-related genetic variations in cfDNA is technically required for the accuracy in ctDNA detection before it can be steadily used in early cancer diagnosis^17^. Thirdly, consistency should be proved. Namely, genetic variations detected from ctDNA and tDNA (tumor DNA from solid tumor cells) should be proved to be cancer-related. Besides, the evaluating indicator(s) for the assessment of consistency should be reliable and practical.

In order to focus on the above questions, our previous studies explored new resin to increase the recovery rate in ctDNA extraction and examined the background somatic mutations originated from blood cells and cfDNA in 821 non-cancer individuals based on ultra-deep sequencing and we also described the baseline reference^18^. Now we are focusing on the understanding of ctDNA profiles in cancer patients after we developed an updated method called “Sec-Seq” with exogenous molecular barcoding and custom-probe capture to suppress the systematic errors and detect the genetic alterations including mutations, insertion and deletion in early stages of lung cancer.

Here we report our studies on consistency of ctDNA and tDNA in 30 patients containing pulmonary space occupying pathological changes according to traditional clinic diagnosis. We detected the ctDNA by ultra-deep sequencing for 50 cancer-associated genes encompassing region coverage of 29 kb, proving that the diagnosis based on circulating tumor DNA provided about 50% and 75% positive detection rate in phase I and II of lung cancer patients, respectively. Our studies showed the Sec-Seq approach explored in our group can further promote the use of ctDNA in early lung cancer diagnosis and personalized lung cancer therapy.

## Methods

### Ethics

This study was approved by the Shenzhen Second People’s Hospital and Peking University Shenzhen Hospital. All the experiments were performed in accordance with guidelines and regulations of the Shenzhen Second People’s Hospital and the Peking University Shenzhen Hospital. Written consent was obtained from each patient and all analyses were performed anonymously.

### Patients

A total 30 patients were included in this study with the clinical pathological information in **Table S1**, which consisted of 2 pulmonary tuberculosis (PTB), 1 organizing pneumonia (OP) and 27 lung cancers including lung adenocarcinoma (LUAD), squamous cell carcinoma (SCC), adenosquamous carcinoma (ASC), sarcoma (SARC) and small cell lung cancer (SCLC). There are three forms including cfDNA, WBC and tissue for each patient. The sample group comprised 33.3% females and 66.7% males. The participant ages ranged from 36 to 77 years, with a median of 59.5 years old. Our analysis of the patient clinical information revealed no significant genetic diversity in the samples. The statistical analysis of sample information is summarized in **Table 1**.

**Table 1.** Clinical features of 30 patients containing pulmonary space occupying pathological changes.

| Characteristic | Parameter value |
| --- | --- |
| **Age, years** |  |
| Mean (SD) | 59.5 (10.8) |
| Median (Range) | 62.5 (36-77) |
| **Sex, n (%)** |  |
| Male | 20 (66.7) |
| Female | 10 (33.3) |
| **Pathological diagnosis, n (%)** |  |
| non-small cell lung cancer (NSCLC) | 26 (86.7) |
| Lung adenocarcinoma (LUAD) | 19 (63.3) |
| squamous cell carcinoma (SCC) | 4 (13.3) |
| small cell lung cancer (SCLC) | 1 (3.3) |
| ASC with SARC | 1 (3.3) |
| ASC | 1 (3.3) |
| SARC | 1 (3.3) |
| pulmonary tuberculosis (PTB) | 2 (6.7) |
| organizing pneumonia (OP) | 1 (3.3) |
| **Tumor stage, n (% in all samples)** |  |
| I A | 7 (23.3) |
| II A | 2 (6.7) |
| II B | 4 (13.3) |
| IIIA | 12 (40.0) |
| IVA | 1 (3.3) |
| NA | 4 (13.3) |
| **Smoking status, n (% in all samples)** |  |
| Non-smoker | 14 (46.7) |
| Smoker | 16 (53.3) |

### Blood plasma isolation

For each sample, 10 ml of blood in a cell-free DNA BCT blood collection tube (Streck, Omaha, US) was collected and then centrifuged at 1600 g for 10 min at 4°C to roughly separate the sample into plasma and blood cells. The upper phase was then transferred into a new tube, leaving around 3-4 mm of “buffering” layer from the buffy coat after the centrifugation and avoid contaminating the plasma layer by blood or cell debris. The plasma was further centrifuged at 16000 g for another 10 min to remove the cell debris. The upper clear layer was then aliquoted into 2-ml tubes, clearly labelled with the patients’ identity and immediately stored at −80°C for cfDNA extraction.

### Extraction of cfDNA

Each extraction of cfDNA was performed from 3 ml of plasma. Extraction of cfDNA was conducted using the QIAamp Circulating Nucleic Aid Kit (Qiagen, Hilden, Germany). The concentration of extracted DNA was measured using the Qubit 3.0, dsDNA high-sensitivity assay (Life Technologies, Carlsbad, CA). All methods were performed according the manufacturers’ instructions.

### Spike in control

The sensitivity and precision of the current method were evaluated by Horizon’s Partners Spike-in control (Horizon, Cambridge, UK) using our custom-designed probes from <u>Integrated DNA Technologies (</u>IDT). Briefly, we performed a serial dilution (from 0.0005 to 1) using the wild-type reference genome and the provided reference standard. We performed multiplex PCR, and then library construction and sequenced on the HiseqX10 (Illumina, CA, USA). Six reference variants were included in our primer set: EGFR (L858R, T790M), KRAS (G12D), NRAS (Q61K, A59T) and PIK3CA (E545K). All other primers were used as the background.

### Extraction of genomic DNA from WBC

Genomic DNA of WBCs was extracted by the Qiagen DNA mini kit (Qiagen, Hilden, Germany). The concentration of extracted DNA was measured using the Qubit 3.0, dsDNA high-sensitivity assay (Life Technologies, Carlsbad, CA). All methods were performed according the manufacturers’ instructions.

### Extraction of genomic DNA from FFPE

Each extraction of genomic DNA was performed from FFPE. Extraction of genomic DNA was conducted using the QIAamp DNA FFPE Tissue (QIAGEN, Hilden, Germany). The concentration of extracted DNA was measured using the Qubit 3.0, dsDNA high-sensitivity assay (Life Technologies, Carlsbad, CA). All methods were performed according the manufacturers’ instructions.

### Probe hybridization capture and sequencing library construction

We developed a probe hybridization capture method to enrich cancer-associated genes. Fifty cancer-associated genes, summarized in **Table 2**, were included and covered a 29139 bp region. A total 2751 mutations were defined as hotspot mutations (**Table S2**). We performed libraries on circulating DNA from 3ml plasma using the KAPA kit with standard procedures. We performed libraries on 200ng DNA from WBC or FFPE using the KAPA kit with standard procedures as shown in **Figure 1**. Barcoding was employed to distinguish real biological mutations from asymmetric DNA errors and sequencing erroes. Indexed libraries were constructed, and hybrid selection was performed with a custom xGen Lockdown Probes Library (IDT) in multiplex. The post-capture multiplexed libraries were amplified with Illumina backbone primers for 16 cycles of PCR using 1× KAPA HiFi Hot Start Ready Mix and then sent to WuxiNextCODE on the Illumina Hiseq X10 platform (Illumina, Beghelli, CA, USA).

**Figure 1.**
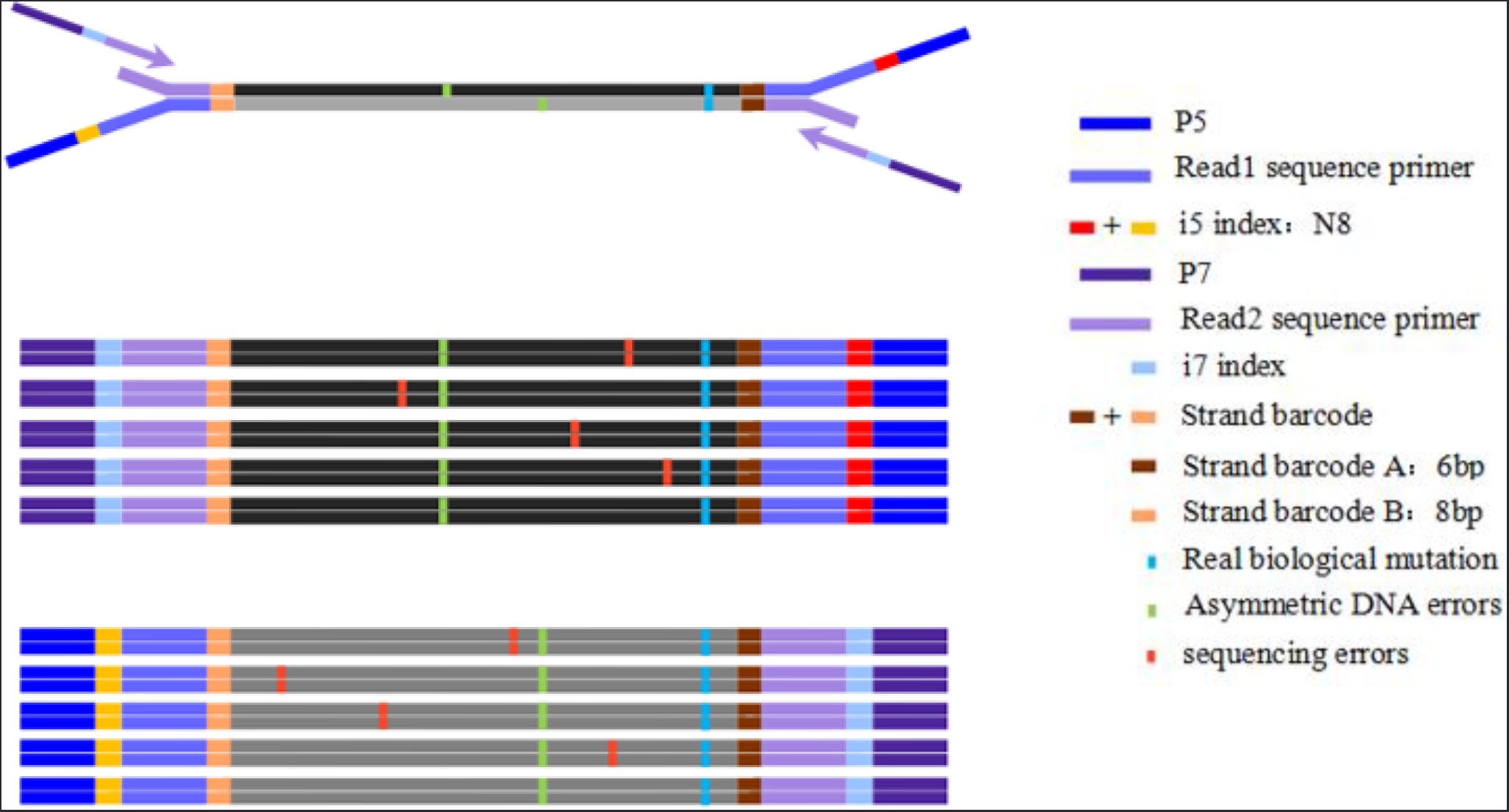
Diagram depicting the use of Systematic error correction in ultra-deep sequencing (Sec-Seq) barcode adapters to suppress errors.

### Data filtering and analysis

We performed ultra-deep target sequencing on 50 cancer-associated genes for both cfDNA and WBC DNA. For each sample, the average sequencing depth was 40000×. The sequencing data was first mapped to the human reference genome (human genome build19; hg19) by Burrows–Wheeler transformation (BWA, Version: 0.7.5a-r405) software package, converted to mpileup format for downstream analysis. We set two filtering criteria to filter the reads: 1) read sequences with mutant allele frequency higher than 5% in a single read were deleted; and 2) bases with base quality lower than 30 were deleted. Loci with sequencing depth less than 10000× were removed for further analysis.

### Sensitivity evaluation

The sensitivity and precision of the current method were evaluated by Horizon’s Partners Spike-in control (Horizon, Cambridge, UK) using our 207-pair primer set. Briefly, we performed a serial dilution (from 0.0005 to 1) using the wild-type reference genome and the provided reference standard. We performed multiplex PCR, and then library construction and sequenced on the HiseqX10 (Illumina, CA, USA). Six reference variants were included in our primer set: EGFR (L858R, T790M), KRAS (G12D), NRAS (Q61K, A59T) and PIK3CA (E545K). All other primers were used as the background.

### Reproducibility evaluation

To validate the reproducibility of our methods, we drew blood from two healthy individuals, spilt each sample in half, and performed library construction and sequencing independently. We also collected FFPE from two patients, spilt each sample in half, and performed library construction and sequencing independently. By comparing the sequencing data of the two replicates in cfDNA or FFPE samples, we were able to evaluate the stability of the methods and remove background noise. Positions with a sequencing depth lower than 10000× and mutant allele frequency below 0.001 were excluded in the correlation study.

### Mutant allele frequency in cfDNA and WBCs in the 30 patients

We analysed the correlation of the mutant allele frequency between cfDNA and genomic DNA of WBCs. The mutant allele frequency was defined as mutant allele frequency divided by total reads covering the locus. For example, for a particular position, the total sequencing depth is 10000; we obtain 9990 for A, 3 for C, 5 for G, and 2 for T, and thus the mutant allele frequency for this position is (3+5+2)/10000=1/1000. Paired cfDNA and WBCs collected from the same patient (at the same time) were analysed. A total of 29 kb nucleotides in 50 genes were covered in this study. We calculated the average mutant allele frequency of each position in 30 patients in cfDNA and WBCs. We removed the positions with mutant allele frequency higher than 0.1.

### Mutant allele frequency in cfDNA and tDNA in the 30 patients

We analysed the correlation of the mutant allele frequency between cfDNA and genomic DNA of solid tumor tissue (tDNA). The mutant allele frequency was defined as mutant allele frequency divided by total reads covering the locus. For example, for a particular position, the total sequencing depth is 10000; we obtain 9990 for A, 3 for C, 5 for G, and 2 for T, and thus the mutant allele frequency for this position is (3+5+2)/10000=1/1000. Paired cfDNA and tDNA collected from the same patient were analysed. A total of 29 kb nucleotides in 50 genes were covered in this study. We calculated the average mutant allele frequency of each position in 30 patients in cfDNA and tDNA. We removed the positions with mutant allele frequency higher than 0.1.

## Results

### Validating the sequencing coverage of plasma cfDNA

The length distribution over all sequences of plasma ctDNA largely fit the normal distribution, with most lengths between 60 to 160 bp. Therefore, the sequences in this range of lengths were selected for further analysis (**Figure 2A**). GC content across all bases of plasma cfDNA was between 45%-55%. The region of reads between 1 to 20 bp fluctuated widely and was therefore removed during quality control (**Figure 2B**). Evaluation of each amplicon of all plasma samples showed the depth of most amplicons was over 20,000-fold (**Figure 2C**). Saturation curve suggested the average templates for each plasma sample were around 1800 and dataset of 6G was sufficient to capture all the information provided in the templates (**Figure 2D**). Baseline noises of conventional NGS and Sec-Seq after sequencing error corrected are indicated at each base in the captured regions of interest. The noise is lowered by more than 3 orders of magnitude in Sec-Seq (**Figure 2E**). The quality score of every base called during the sequencing runs illustrated the sequencing accuracy was reliable (**Figure S1 A)**. We also did the validation of tDNA in tumor tissue and validation of gDNA in white blood cells in the initial process of data analysis (**Figure S1 B-I**).

**Figure 2.**
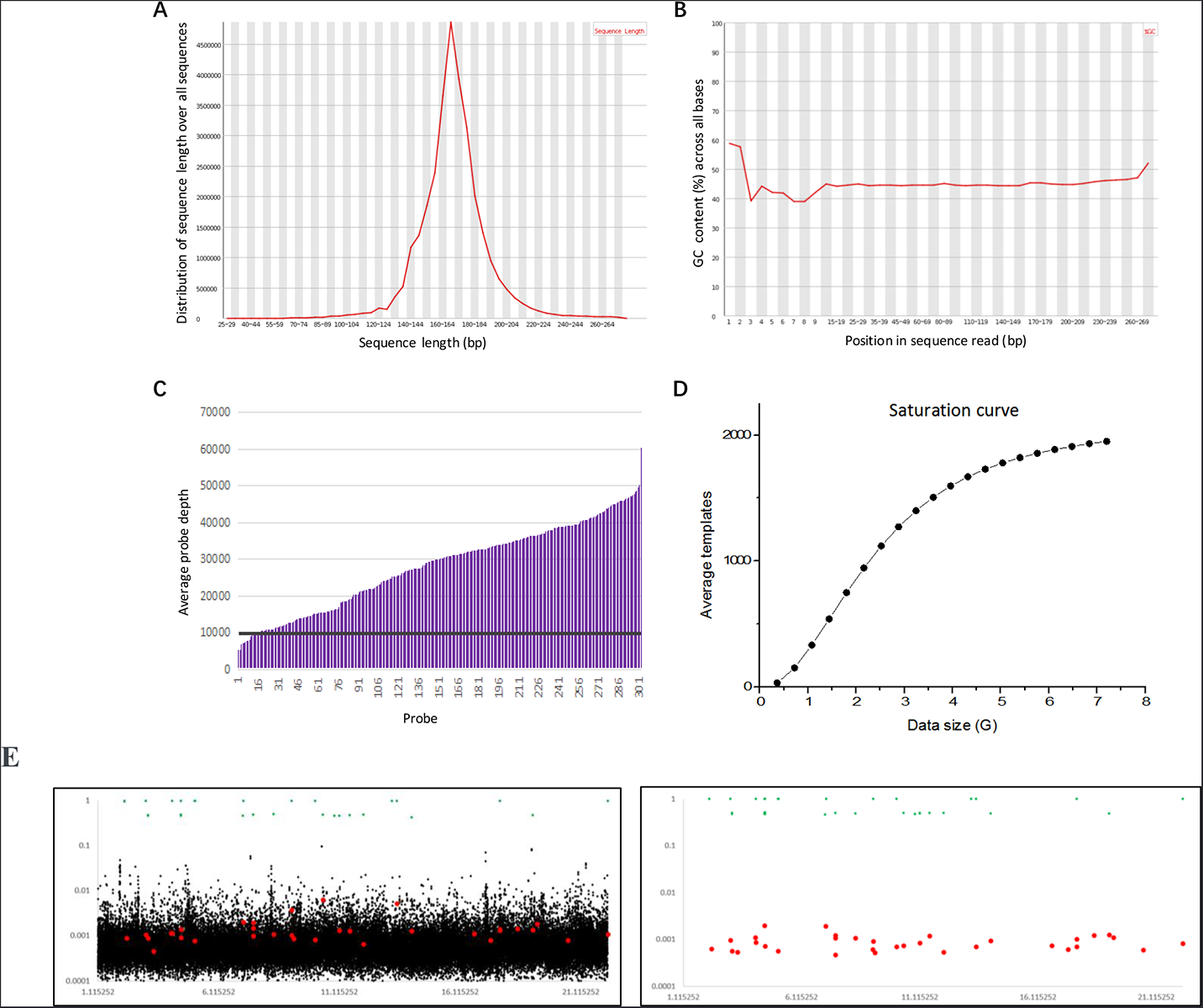
Sequencing coverage of cfDNA. **A**) Distribution of sequence lengths over all sequences of plasma cfDNA. The sequence length distribution shows most lengths are between 80 to 240 bp, which were selected for further analysis; **B**) GC content across all bases of plasma cfDNA. The region of reads between 1 to 15 bp fluctuates widely and was therefore removed during quality control; **C**) The depth of each amplicon of all plasma samples, where the depth of most amplicons is over 20,000x; (**D**) The saturation curve shows the average templates were around 1500-2000 with average data set as 5G; (E) Sec-Seq error correction.

### Validating the detecting sensitivity using the standard reference

Horizon Standard Reference libraries covering six known tumor-specific hotspot mutations in EGFR, NRAS, PIK3CA and KRAS genes were used to evaluate the sensitivity of the Sec-Seq approach used in cfDNA analysis. The average size of the reference genome is around 160 bp, mimicking the plasma cfDNA that we derived from blood plasma. We examined the sensitivity and precision of this cfDNA reference standard using our sample and library preparation method then tested the sensitivity of the method at variant allele frequencies of 0.0005, 0.001, 0.005 and 0.01. Libraries with eight exogenous barcodes were sequenced at an average depth of >20000-fold coverage before the allele frequencies detected for six reference mutation sites were calculated. The results demonstrated that our method could detect more than 90% of variant alleles at a frequency of 0.001 with acceptable fluctuation and the stochastic fluctuation was significantly reduced at the reference 0.005 and 0.01 (**Figure 3**). Therefore, threshold of average allele frequency for cfDNA and tDNA detection was subsequently defined as 0.001, and threshold of average allele frequency for quantitative detection in tumor tissue was defined as 0.02.

**Figure 3.**
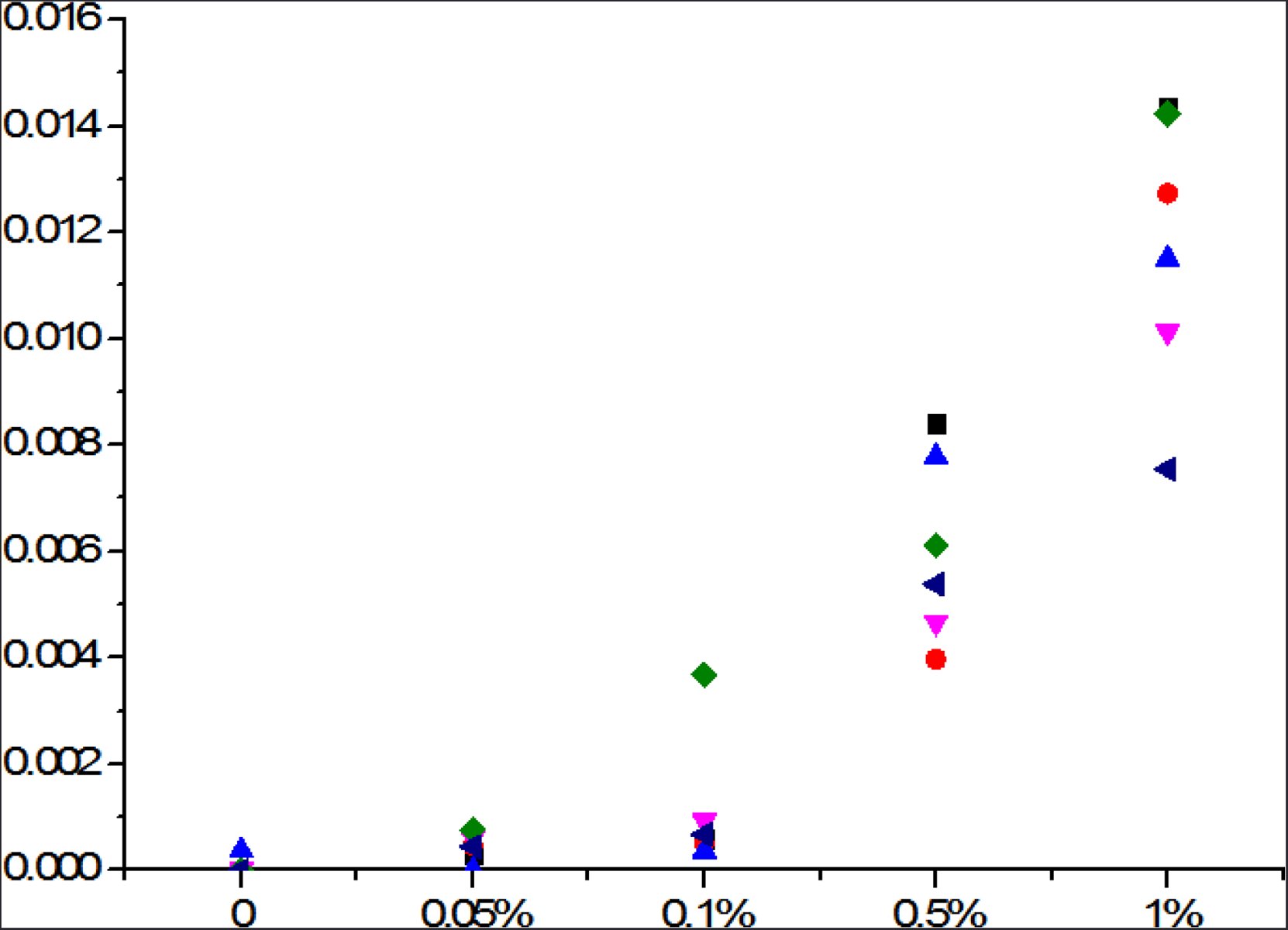
Sensitivity test using Horizon Reference Standard for cfDNA. The reference was manufactured from engineered human cancer cell lines with an allele frequency of 0%, 0.05%, 0.1%, 0.5% and 1.0%. Deep sequencing was performed and the allele frequencies detected for six reference mutation sites were calculated.

### Reproducibility

We compared the sequencing information of two technical replicate cfDNA samples (left) from the same healthy individual (two individuals in this study) and two technical replicate FFPE samples (right) from the same lung cancer patient (two patients in this study) to evaluate the reproducibility of the Sec-Seq approach in order to distinguish the true biological alterations from systematic errors. Only a total sequencing depth larger than 10000× and mutant allele frequency larger than 0.1% for cfDNA samples were included in the correlation study. Our data showed that the mutant allele frequency in the cfDNA samples were highly correlated (Adj R^2^=0.8799 for individual 1 and Adj R^2^=0.9589 for individual 2), implying the reproducibility in plasma ctDNA detection of cancer patients will be even better considering the cfDNA concentration is lower in healthy individuals than in cancer patients (**Figure 4**). The mutant allele frequency in the two FFPE samples also showed good correlations (Adj R^2^=0.9354 for patient 1 and Adj R^2^=0.9448 for patient 2) (**Figure 4**), suggesting the Sec-Seq approach can generate satisfactory reproducibilities for all the sample forms in our study (data for white blood cell gDNA not shown).

**Figure 4.**
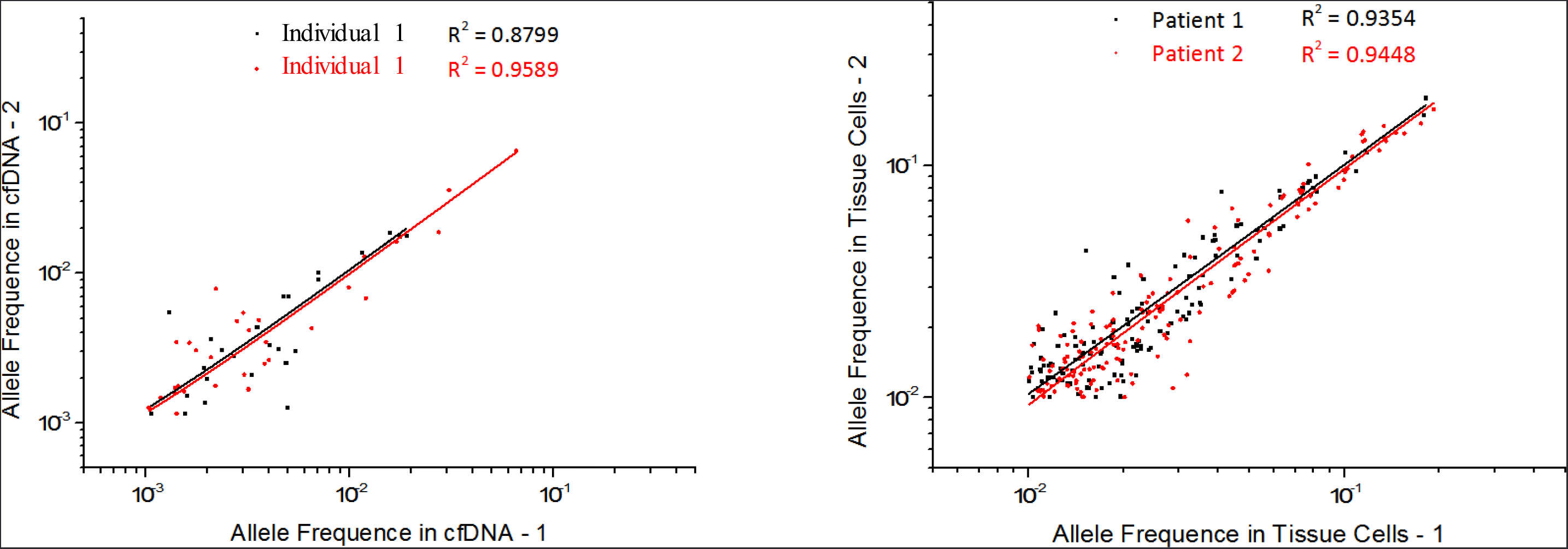
Reproducibility validation in cfDNA and tDNA. We compared the sequencing information of two replicate cfDNA samples (left) and two replicate tDNA samples (right) to evaluate the reproducibility of the Sec-Seq approach. Only a total sequencing depth larger than 10000× and mutant allele frequency larger than 0.1% were included in the correlation study.

### Consistency between pathology diagnosis and plasma ctDNA mutations detected

We collected the pre-operation blood and post-operation FFPE samples from 30 patients containing pulmonary space occupying pathological changes according to our experimental design. By using the Sec-Seq approach, we performed ultra-deep target sequencing on 50 cancer-associated genes (**Table 2**) for plasma DNA (cfDNA) and deep target sequencing for tumor tissue ctDNA, while the white blood cell DNA was also analysed for the somatic mutations in data filtering. Finally, the sequencing data of tumor tissue gDNA for 4 patients were not available The clinic diagnosis was not referred until all the sequencing and technical bioinformatic analysis were completed. The 30 patients turned out to be composed of 2 pulmonary tuberculosis (PTB), 1 organizing pneumonia (OP) and 27 lung cancers diagnosed at stage I to IV (**Table 1** and **Table S3**). We found that the concentration of cfDNA in plasma from cancer patients slightly increased with the progression of cancer, showing average of 20 ng/ml for stage III patients versus 18 ng/ml for stage I patients (**Table S3**), while the trend was not observed for the mutant allele frequency (**Table S3** and **S4**). The ctDNA detection and the diagnosis of SOL of the lung cancer patients demonstrated good consistency between pathology diagnosis and plasma mutations detected. The 3 benign lesion were detected as negative. The overall concordance was 92% (***p <*** 0.001, SPSS Statistics version 19). For all the 23 lung cancer patients with qualified sequencing results of ctDNA detection, 85, 80, 100 and 90% lung cancer patients in stage I, II, III and I-IV, respectively, were predicted to be positive (**Table S5**).

**Table 2.**
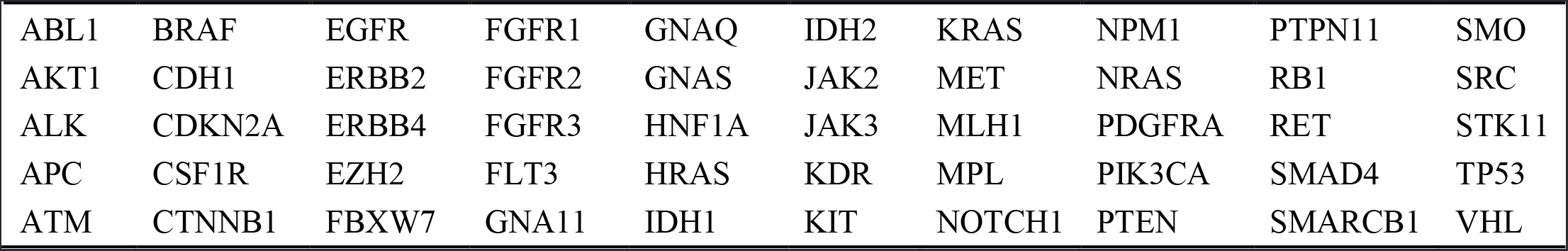
Genes covered by targeted NGS Panel.

### Comparison of alterations in plasma with those in matched tumor tissue for patients with lung cancer

The patient characteristics and gene mutations in matched tDNA and plasma ctDNA sample pairs were summarized while patients were categorized based on stage, age, sex, and pathological diagnosis (**Table 3** and **Figure 5**). Only top nine genes were shown here for simplification (for detailed information of all the 50 genes in the panel, refer **Table S4**). Although the sample capacity was limited, it was still found that the consistency in males was better than in females in stage I lung cancer, which may imply the difference of biological situations or responsive abilities between two genders especially in the ultra-early or early stages of cancers as females usually have a higher level of complexity. The consistency rate was evaluated by the percentage of lung cancer patients having one or more shared variants in tumor tissue and plasma (indicated with red star in **Figure 5**), namely the ratio of the number of patients with ctDNA alterations to the number of patients with the identical alterations in tDNA, which was 78.95% for the overall I-IV stages (15/19 in **Table S5**), and 50, 75, 100 or 100% for patients in stage I, II, III or IV, respectively.

**Table 3.** Consistency analysis of mutations in tDNA and ctDNA.

| Patient no. | Gene | Mutation type | AA mutation | % mutation in tDNA (reads) | % mutation in pre-op ctDNA (reads) | % mutation in WBC (reads) | Detected in tDNA/ctDNA | Tumor stage | Age | Smoking status | Pathological diagnosis |
| --- | --- | --- | --- | --- | --- | --- | --- | --- | --- | --- | --- |
| 1 | IDH1 | SNV | p.R132H | 0.037(5400) | 0.157(1271) | 0.047(10514) | N/Y | II B | 69 | Y | SCC |
|  | TP53 | SNV | p.R273H | 17.8(4626) | 0.118(843) | 0.033(9031) | Y/Y |  |  |  |  |
| 2 | AKT1 | SNV | p.E17K | 0.031(9583) | 0.104(1909) | 0(9433) | N/Y | I A2 | 43 | N | LUAD |
|  | GNAS | SNV | p.R201H | 0.131(9127) | 0.286(1397) | 0.063(11041) | N/Y |  |  |  |  |
| 3 | TP53 | SNV | p.V216M | 19.5(5397) | 0.183(2742) | 0.076(10409) | Y/Y | III A | 74 | Y | (ASC with SARC) NSCLC |
| 5 | KRAS | SNV | p.G12V | 26.7(7783) | 0.126(793) | 0.032(12253) | Y/Y | I A3 | 53 | Y | LUAD |
|  | TP53 | SNV | p.R196Q | 0.108(3689) | 0.232(861) | 0(12789) | N/Y |  |  |  |  |
| 6 | EGFR | INDEL | p.L747_P753>S | 28.9(1212) | 1.32(1806) | 0(16633) | Y/Y | III A | 42 | N | LUAD |
|  | TP53 | SNV | p.Q136* | 20.3(531) | 0.792(1640) | 0.014(13342) | Y/Y |  |  |  |  |
| 7 | IDH1 | SNV | p.R132H | 0.014(7138) | 0.161(1855) | 0.059(6682) | N/Y | III | 48 | N | LUAD |
|  | EGFR | SNV | p.L861R | 0.038(10434) | 0.296(1687) | 0.039(7670) | N/Y |  |  |  |  |
|  | NRAS | SNV | p.Q61R | 25.7(4929) | 0(1103) | 0.528(3783) | Y/N |  |  |  |  |
| 8 | KRAS | SNV | p.G13G | 0.088(10141) | 0.128(1561) | 0.052(19104) | N/Y | I A2 | 42 | Y | LUAD |
| 9 | EGFR | INDEL | p.L747_P753>S | 41.8(17443) | 0(3064) | 0(6572) | Y/N | I A2 | 67 | N | LUAD |
| 10 | EGFR | INDEL | p.E746_S752>A | 16.7(8544) | 0(1844) | 0(8087) | Y/N | I A | 65 | N | LUAD |
|  | EGFR | SNV | p.S752F | 18.2(7892) | 0.105(1898) | 0.012(7740) | Y/Y |  |  |  |  |
| 11 | EGFR | SNV | p.D761N | 23.1(4214) | 0.402(1738) | 0.031(9602) | Y/Y | III A | 63 | Y | SCC |
|  | TP53 | SNV | p.R273H | 41.7(5756) | 2.588(1275) | 0.011(8958) | Y/Y |  |  |  |  |
|  | AKT1 | SNV | p.E17K | 43.4(5247) | 0.125(1598) | 0.088(7920) | Y/Y |  |  |  |  |
| 12 | EGFR | SNV | p.L858R | 40.6(14018) | 0.683(1609) |  | Y/Y | III A | 65 | Y | LUAD |
|  | TP53 | SNV | p.Q192* | 0.214(5137) | 0.162(1234) |  | N/Y |  |  |  |  |
| 13 | TP53 | SNV | p.V73L | 1.946(3236) | 0.243(2468) |  | Y/Y | IV A | 70 | N | LUAD |
|  | FGFR3 | INDEL | p.S779fs*>28 | 0.125(5588) | 0.302(2977) |  | N/Y |  |  |  |  |
|  | IDH1 | SNV | p.R132H | 0.092(8686) | 0.117(3418) |  | N/Y |  |  |  |  |
| 14 | KDR | SNV | p.G873R | 18.1(6200) | 0.638(1410) | 0.020(4986) | Y/Y | II B | 63 | N | (ASC) NSCLC |
|  | TP53 | SNV | p.R196Q | 0.234(7688) | 0.302(1324) | 0.147(4730) | N/Y |  |  |  |  |
| 15 | KRAS | SNV | p.G12A | 29.1(5745) | 0(3353) |  | Y/N | II B | 69 | Y | LUAD |
| 16 | IDH1 | SNP | p.G105G | 0(3335) | 0.140(1419) | 0(410) | N/Y | III A | 67 | Y | SCC |
| 17 | SMARCB1 | SNV | p.? | 11.3(10082) | 31.4(1517) |  | Y/Y | III A | 58 | Y | SCLC |
|  | PDGFRA | SNP | p.V824V | 14.5(5400) | 35.6(1754) |  | Y/Y |  |  |  |  |
|  | TP53 | NA |  | 0.740404457 | 0.423145401 |  |  |  |  |  |  |
| 18 | TP53 | SNV | p.M237T | 0.068(1461) | 0.133(2988) | 0.028(7075) | N/N | N/A | 65 | Y | OP |
| 19 | EGFR | SNV | p.L858R | 10.03(13817) | 0.846(2362) | 0.040(14706) | Y/Y | III A | 55 | N | PC, (SARC), (NSCLC) |
|  | TP53 | SNV | p.R110L | 2.68(5889) | 0.375(2133) | 0.046(12941) | Y/Y |  |  |  |  |
| 21 | TP53 | SNV | p.G245S | 0.050(7983) | 0.113(1758) | 0.0872(18352) | N/Y | I A | 66 | N | LUAD |
|  | EGFR | SNV | p.L858R | 20.3(14244) | 0(2183) | 0.017(22533) | Y/N |  |  |  |  |
| 22 | PIK3CA | SNV | p.H1047R | 0.091(10908) | 3.73(2410) | 0.037(29389) | N/Y | III A | 61 | Y | SCC |
|  | TP53 | SNV | p.? | 0.072(12413) | 0.622(2570) | 0.032(27385) | N/Y |  |  |  |  |
| 23 | KRAS | SNV | p.G12C | 4.66(9002) | 0(1738) | 0.036(16247) | Y/N | II A | 77 | Y | LUAD |
| 24 | TP53 | SNV | p.V73L | 1.40(2067) | 0.213(2814) |  | N/Y | N/A | 55 | Y | PTB |
| 25 | TP53 | SNV | p.R202C | 0.120(2486) | 0.102(1952) | 0.079(17507) | N/Y | III A | 66 | N | LUAD |
| 26 | TP53 | SNV | p.R282W | 0.106(10365) | 0.221(905) | 0.005(17210) | N/Y | II A | 45 | Y | LUAD |
| 27 | JAK3 | SNV | p.? | 0.039(7609) | 0.108(1965) | 0.028(14164) | N/Y | III A | 36 | Y | LUAD |
| 30 | TP53 | SNV | p.Y234H | 0.066(2986) | 0.179(1672) |  | N/Y | N/A | 77 | N | OP |
|  | EGFR | SNV | p.G719A | 8.00(6735) | 0(1849) |  | Y/N |  |  |  |  |

**Figure 5.**
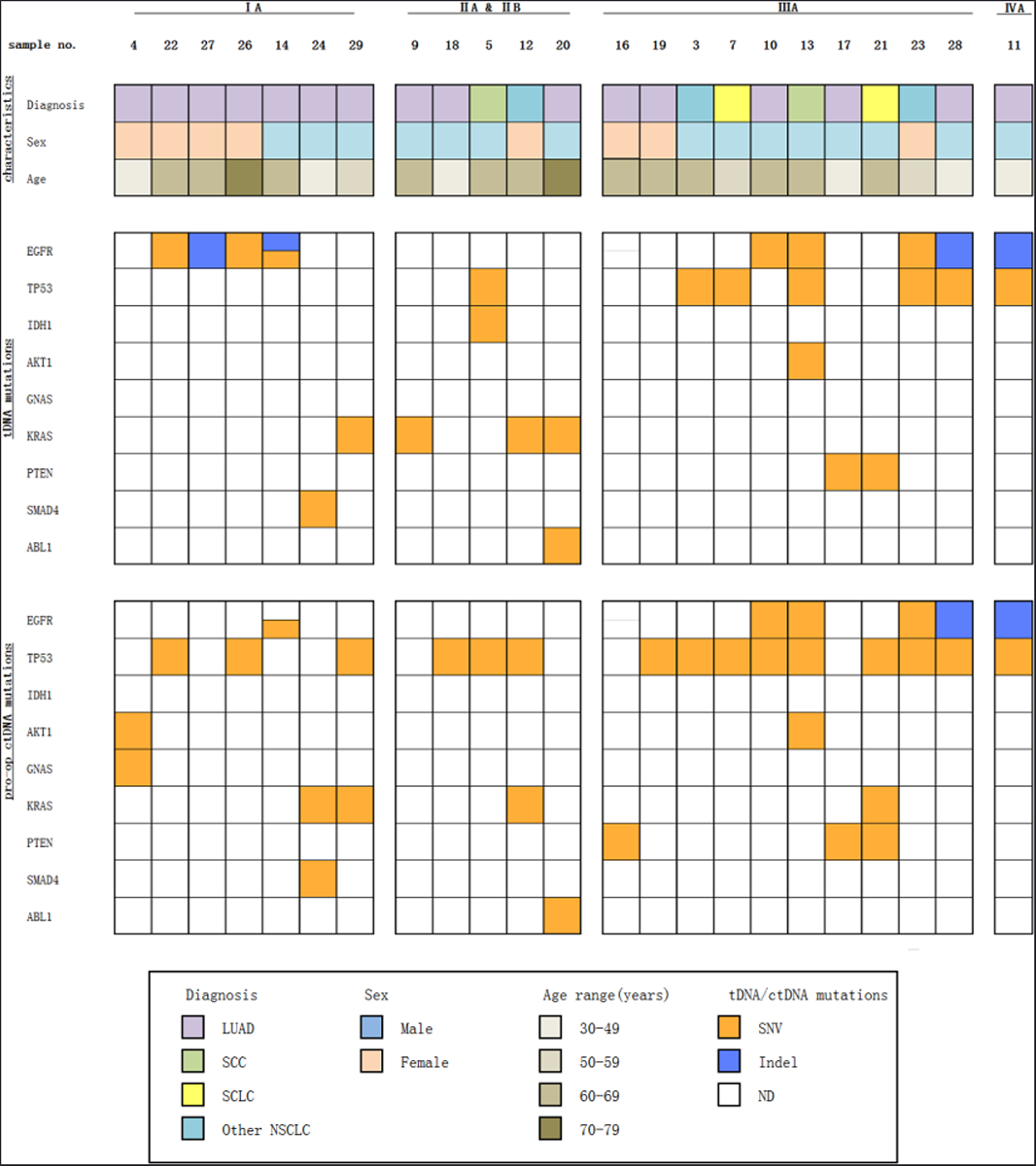
Consistency analysis of mutations in tDNA and ctDNA. Patients were categorized based on stage, age, sex and pathological diagnosis (top); type of mutation per gene in tDNA (middle) and pro-op plasma ctDNA (bottom). tDNA and plasma ctDNA samples with one or more mutations in the same gene are indicated. Orange indicates SNV, blue indicates Indel, and white indicates no genetic variations were detected in that gene in tDNA (middle) or pro-op plasma ctDNA (bottom). Only top nine genes in the panel were shown here for simplification. The consistency rate was evaluated by the percentage of lung cancer patients having one or more shared variants in tumor tissue and plasma.

### Comparison of ctDNA detection and tumor biomarkers in plasma from patients with lung cancer

Most of the pre-surgery plasma samples were analyzed for the presence of the following tumor biomarkers including CA125, CA153, CEA, NSE, CA19-9, CYFRA21-1, SCC (squamous cell carcinoma antigen) and AFP (**Figure 6** and **Table S6**) while only the former six were taken into consideration for comparison with plasma ctDNA for the detection of latter two failed in some patients. Finally, there were 22 patients in this analysis. The positive detection was defined as one or more of the above six biomarkers detected as positive. The overall positive rate of plasma ctDNA was 84% while that of biomarkers was 35%. More notably, the advantage and significance of plasma ctDNA detection is reflected in early stages in lung cancer. It is obvious that the positive rate of plasma ctDNA for stage I and II was higher than 70%, when tumor biomarkers were insensitive.

**Figure 6.**
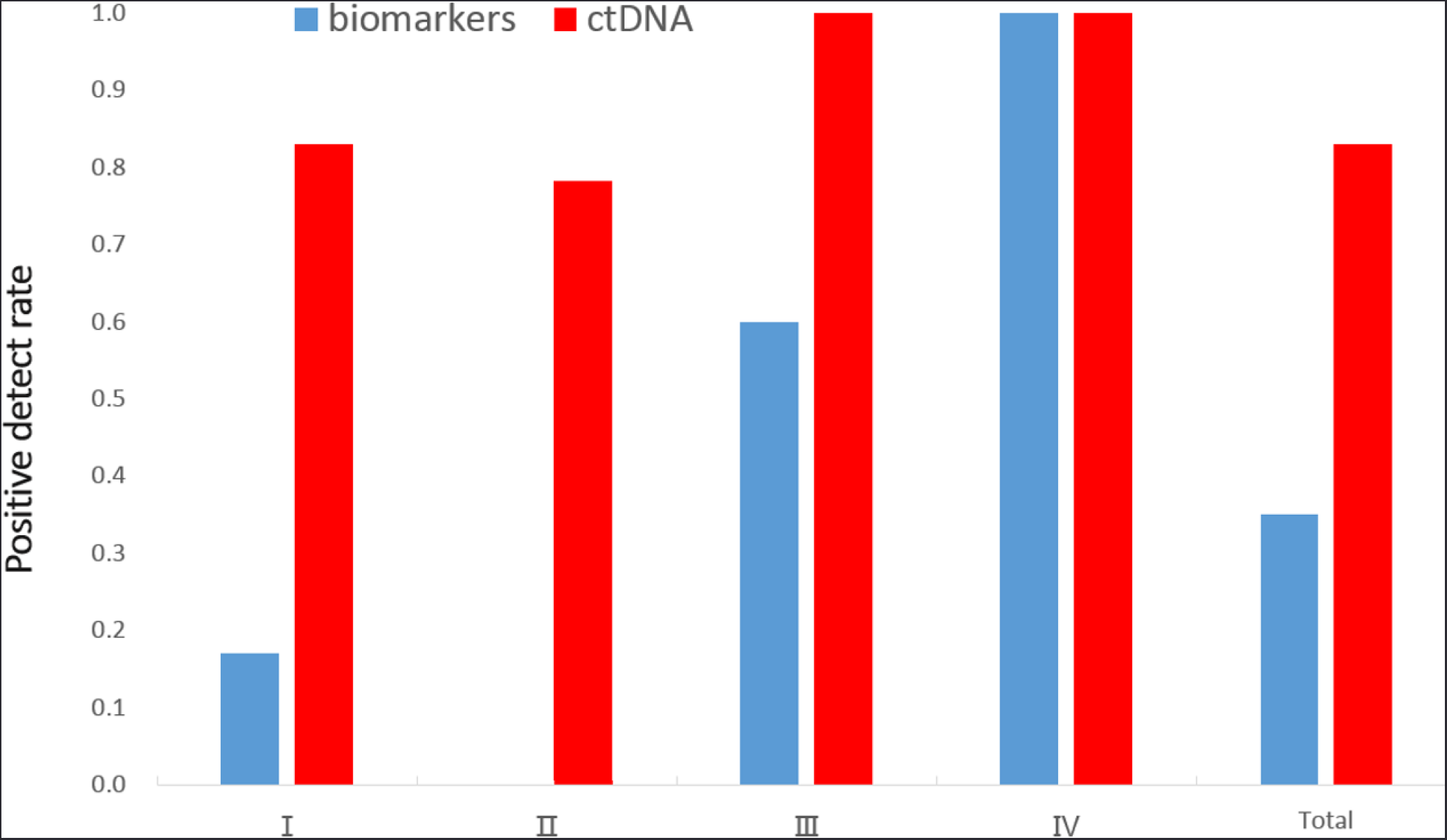
Comparison of positive detection rates of plasma ctDNA and tumor biomarkers. The pre-surgery plasma samples were analyzed for the presence of the following tumor biomarkers including CA125, CA153, CEA, NSE, CA19-9, CYFRA21-1. The positive detection was defined as one or more of these six biomarkers detected as positive.

## Discussion

The ctDNA is released to the peripheral blood in the initiation stage of tumorogenesis, meaning theoretically plasma ctDNA can be detected earlier than CT or other traditional clinical methods in cancer diagnosis. Therefore, cfDNA and ctDNA detection has emerged as the research and commercial frontier of non-invasive cancer biomarkers for monitoring and treatment of cancer^19-24^ in recent years. cfDNA may not 100% represent the condition of tumor progress, but it is undoubtedly a powerful tool in clinical practice to help diagnose the cancer and evaluate treatment effect to some extent, particularly when the tumor tissue biopsy is hardly feasible. Undoubtedly, the ambition of plasma ctDNA is to promote its use in early diagnosis for cancers with the technical development of NGS and cost decrease of sequencing.

Here we developed a method called Sec-Seq aimed to facilitate the systematic errors correction in ctDNA detection and proved its effectiveness in positive detection of lung cancers, which preliminarily indicates the advantages of plasma ctDNA in diagnosis of early-stage cancers. As a novel probe hybridization capture sequencing technology, Sec-Seq allows ultrasensitive and direct evaluation of sequence changes in circulating cell-free DNA. Although we used it in experiments designed for lung cancer diagnosis, it can be used for any other cancer type.

We previous reported that somatic mutations in the blood cells significantly contribute to mutations in cfDNA in healthy individuals^18^. Now we still noticed this in our current studies (**Table S4**). These results emphasize again the importance of sequencing both cfDNA and blood cells to remove the background mutations contributed by blood cells.

In our studies, using the Sec-Seq approach, the ctDNA mutations in plasma samples obtained before surgery had a high concordance rate to mutations found in primary tumor tissue, which showed the superiority compared with the detection of tumor markers expression. Besides, the Sec-Seq for cfDNA assay had a satisfactory specificity, sensitivity, and PPV for early diagnosis for patient with SOL of the lung, while these results need to be further confirmed in larger size of samples. Notably, no tumor-derived alterations were identified in the plasma of the patient with benign lesion of the lung in our study.

Our data showed the overall genetic variations (**Figure S2**) has the comparable concordance between ctDNA and tDNA, most of which are of passenger genes. We believe that more data are required for the elucidation of tumor-related genetic alterations, as the COSMIC hotspots are mainly aimed to target the driver genes.

We also observed the unconsistency between ctDNA and tDNA in our studies (**Figure 5**) and analysed ranking of the top 15 genes in our panel in terms of variation probability in patients (**Figure 7**). It was found that TP53 is the most mutated gene in both ctDNA and tDNA, which has been reported by previous studies. Besides, mutations in TP53, EGFR, PTEN, RET, PIK3CA, KRAS, KDR and ATM were detected in both plasma and tumor tissues in most patients, while the frequencies are partially consistent. We think the cancer heterogeneity of tissue and space-time of ctDNA may cause the differences. The unconsistency may not means the low sensitivity or false positive result in ctDNA detection, but be originated from the biological reasons including the secretory characteristics, metabolic half-life and cancer heterogeneity. We propose different panels should be considered for DNA biomarkers of tumor tissue and plasma cfDNA, since they apparently do not share the ranking of genes in terms of detectable variations.

**Figure 7.**
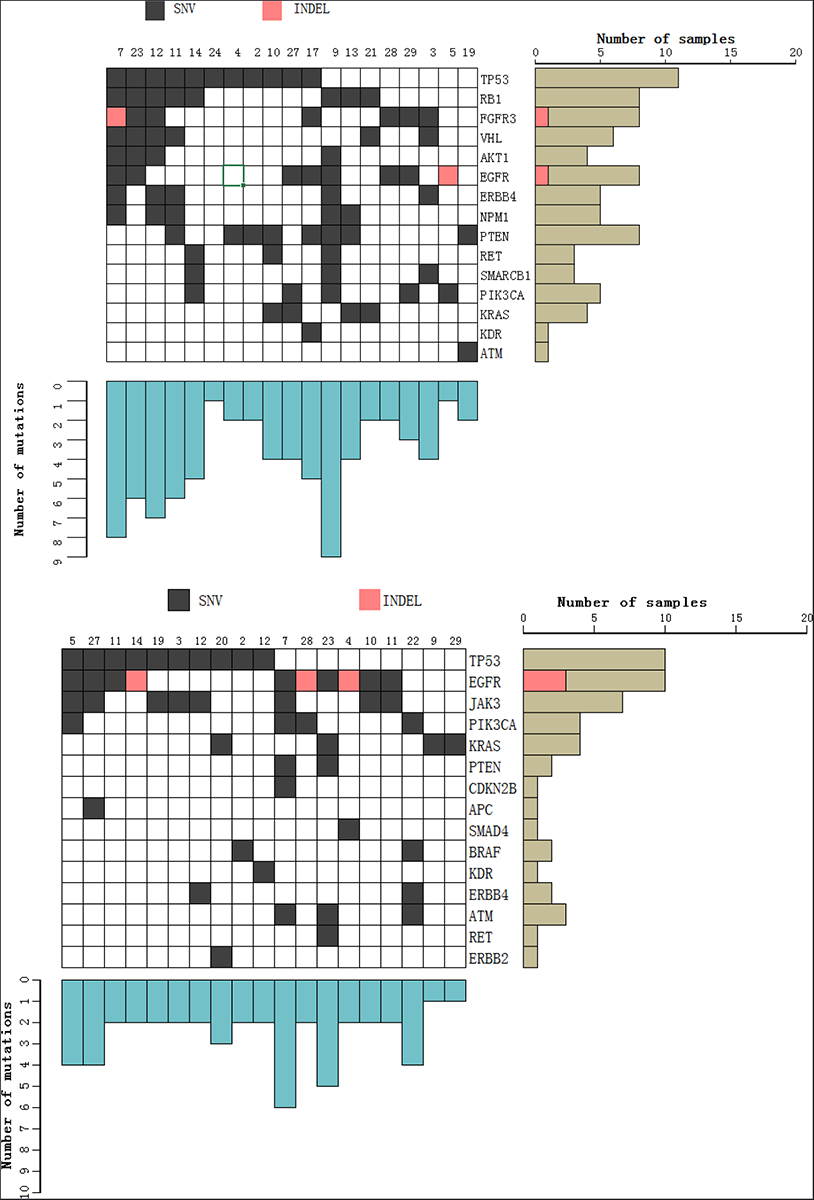
Mutant allele frequency of all the top genetic variations in ctDNA (upper) and tDNA (lower)

One more difficulty needs to be pointed out for tDNA mutation detection. We collected the matched FFPE samples but some failed in the DNA extraction or sequencing, resulting in no significant data for 4 lung cancer patients with the reason(s) not determined, which implies the quality control of FFPE sample is very important considering its complexity.

Lastly, we are focusing on a larger sample size to refine the baseline spectrum and more efficient bioinformatics pipeline to increase the accuracy and sensitivity in the mutation detection of ctDNA. More results of our Sec-Seq approach to establish early cancer diagnosis strategies in cancers will be published in upcoming articles.

## Supplementary data

Supplementary Figure S1 Sequencing coverage of ctDNA, tDNA or white blood cell gDNA

Supplementary Figure S2 Landscape of genetic alterations in lung cancer patients

Supplementary Table S1 Summary of clinical pathological information

Supplementary Table S2 50-gene panel and probe information

Supplementary Table S3 Relationship between cfDNA centration and stages of lung cancer

Supplementary Table S4 Sequencing data of the 30 patients

Supplementary Table S5 ctDNA detection and the diagnosis of SOL of the lung cancer patients

Supplementary Table S6 Detection of plasma protein biomarkers

## Acknowledgements

The authors thank Shanghai Mingma for the sequencing services. This work was supported by Shenzhen Innovation Fund of China (Grant No: CKCY2016082916544973); State Key Research Program of China (Grant No: 2016YFA0501604); Shenzhen Technological Innovation Research Program of China (Grant No: JSGG20160428090301587); the National Natural Science Foundation of China (Grant No: 31200563); Shenzhen Basic Research Program of China (Grant No: JCYJ20140819153305695); the Young Scientist Innovation Team Project of Hubei Colleges (Grant No: T201510); the Key Project of Health and Family Planning Commission of Hubei Province (Grant No: WJ2017Z023).

## Author Contributions

J.H., X.L. and C.L. designed the study. Y.X, F.C., X.W. and G.T. collected original blood samples and tissue samples from the patients. F.X., D.Y. and X.Y. carried out the experiments. X.T. and S.L. summarized and performed the statistical analysis including the clinical information of all the patients. X. L. wrote and revised the manuscript. All authors read and approved the final manuscript.

